# Multidomain triple target capture for navigating complex symbioses

**DOI:** 10.64898/2026.06.18.733235

**Authors:** Pranoti Giram, Shamim Ahmed, Dexcem J. Pantinople, Heather R. Jordan, Ryan A. Folk

## Abstract

Amplicon sequencing remains the benchmark technique for characterizing microbial communities, but the limitations of PCR bias and single-locus targets conspire to limit conclusions. Metagenomics, the primary alternative, shares with amplicon sequencing challenges of economic scaling at adequate sequencing depth. Targeted enrichment strategies can improve the data resolution and economics of sequencing efforts while reducing methodological bias. Here, we develop a target capture strategy for metagenomic characterization of eukaryotic ribosomal DNA and root nodule symbiosis genes and test it in the metagenomes of plant root nodules. We utilize biotinylated RNA probes to selectively capture genomic regions of interest from complex environmental DNA samples, avoiding forms of PCR bias that can undermine community characterization and overcoming the need for conserved priming sites often lacking in functional genes. We designed custom probe sets targeting conserved flanking regions of eukaryotic ITS and known root nodule symbiosis (RNS)-related genes and tested them on diverse root nodule metagenomes and a mock community. We observed high recovery of target loci from samples, very high on-target read proportions, and enhanced detection of low-abundance taxa compared to amplicon sequencing, including stronger alignment with known mock community compositions. This approach will enable deeper insights into the phylogenetic diversity of eukaryotic symbionts, their genomic adaptations, and the functional potential of symbiotic interactions in a cost-effective manner suitable for large-scale projects. This strategy advances our understanding of microbial community dynamics and symbiotic relationships in natural and anthropogenic ecosystems.

## INTRODUCTION

The structure of bacterial and fungal communities is generally studied using two methods, namely (1) amplicon sequencing and (2) shotgun sequencing (Edwin et al., 2025). Amplicon sequencing of bacterial 16S rRNA genes and fungal ITS regions is widely used because it is relatively cost-effective, sensitive, and sequences can be queried against taxonomically expansive databases like SILVA (Quast et al., 2013), GreenGenes (DeSantis et al., 2006), and UNITE (Nilsson et al., 2019). Yet its broad use has led to a general acknowledgment of the biases and limitations of short amplified fragments. Amplicon sequencing is limited by low taxonomic resolution enabled by querying short regions, PCR primer bias due to the reliance on priming sites, and poor performance on the degraded DNA often seen in environmental samples (Abellan-Schneyder et al., 2021). PCR primer bias is particularly limiting, because it generally prevents application to functional genes (which tend to have few conserved sites appropriate for priming) and even for conserved rDNA priming sites are sometimes problematic and primer universality is less than desired (Abellan-Schneyder et al., 2021), leading to a biased assessment of diversity (Rathod & Silverman, 2026; but see Tessler et al., 2017). In general, amplicon sequencing can over- or underestimate taxa, miss rare but important taxa, and is unable to directly generate functional information, which is more commonly achieved indirectly via functional annotations of taxonomic IDs (Gasc & Peyret, 2018).

Shotgun sequencing addresses various shortcomings of amplicon sequencing; it provides high resolution at both taxonomic and functional levels (Hillmann et al., 2018), detects novel genes, works well on degraded samples, is less biased in terms of relative abundance and taxon dropout, and generally detects more microbial diversity than 16S rRNA gene sequencing (Madison et al., 2023). The chief shortcoming of shotgun sequencing is that it is expensive. Low pass sequencing cannot generally be used on community data effectively because diverse communities require deep sequencing to reliably detect rare taxa, and host DNA often dominates samples from various sources such as plants, limiting microbial DNA recovery and also requiring deep sequencing to overcome (Hillmann et al., 2018). Shotgun sequencing is also limited by the availability of high quality references given that whole-genome queries are most appropriate for this data type but the taxonomic coverage of high-quality genome assemblies is far less than 16S and ITS databases, resulting in likely false positive IDs. The alternative of limiting the reference to well-populated loci greatly decreases the economic efficiency of this technique.

Target capture is a cost-effective method (Rohland & Reich, 2012) that helps to overcome the key limitations of both amplicon and shotgun sequencing: it enriches specific genomic regions leading to lower sequencing effort and low costs in what is otherwise essentially a shotgun experiment, and generally avoids PCR primer bias via a different strategy for targeting loci (Mertes et al., 2011). Target capture is generally used in solution biotinylated RNA probes (also DNA in some workflows) to hybridize with and target desired loci for recovery. In essence, target capture is a form of shotgun sequencing that includes an intercalated or subsequent wetlab technique to enrich a subset of the genome in a library preparation. A variety of approaches exist, such as whole-genome hybridization prepared from gDNA (Enk et al., 2014), but most commonly probes are synthesized according to a bioinformatic design. Depending on the commercial solution, target capture can be implemented on standard libraries, such as Illumina and PacBio, allowing it to be compatible with multiple experiments; multiplexed shotgun and target capture experiments have been standard in plant genomics for years (Hale et al., 2020). While probe syntheses are expensive in themselves, they are compatible with high levels of multiplexing, allowing for very low costs for sufficiently large projects (Hale et al., 2020). Relatively low sequencing effort is appropriate for target capture as hybridization efficiency commonly passes 50%, resulting in enrichment of targets hundreds- to thousands-fold; coverage tends to be 100-1000X in many hybridization studies (Weitemier et al., 2014). The other key advantage of target capture is that probe match is quantitative in nature, meaning that assayed DNAs distant from probes tend to yield lower recovery rather than outright failure. Unlike PCR primers, mismatch is generally tolerated (Schott et al., 2017), and this can be up to 30% mismatch under standard reaction conditions, with the hybridization temperature being an experimental variable available to optimize and manipulate this tolerance. A “degenerate” strategy (in this case, meaning representation of loci by probes from multiple taxa) easily allows for broad taxonomic use beyond this tolerance (Johnson et al., 2019), meaning that target capture is naturally suited to reference mismatch.

The literature reviewed thus far indicates that target capture has been used primarily in plants and animals. Target capture should be highly efficient for capturing microbial diversity measurements as it combines attractive features of amplicon sequencing and shotgun sequencing. So far, target capture has seen limited use in microbial ecology, although it is conceptually similar to GeoChip (He et al., 2007) with the key difference of being a sequencing assay with custom flexible functional panels and multiplexing with microbial barcodes. However, a 16S rRNA panel covering the entire region was recently reported using the technique, which was designed by clustering analyses of 16S database sequences and generating tiled probe sequences from cluster centroids (Beaudry et al., 2021). This panel has been shown to increase 16S by more than 400-fold, resulting in high coverage of regions covered by standard microbial databases at reasonable sequencing effort, and is able to generate community profiles similar to amplicon and shotgun sequencing methods (Beaudry et al., 2021). Most importantly, greater application of target capture would benefit from two things generally, (1) a probe panel for eukaryotic microbial DNA, and (2) further empirical assessment of the performance of target capture in comparison to alternative methods.

A third benefit of target capture would be to (3) leverage the flexibility of probe-based capture to assess microbial function through direct capture of functional genetic loci, which would release microbial ecologists from relying on guessing function from taxonomic annotations. Inferring function from taxon IDs is prevalent but subject to strong assumptions as microbial functions often vary at the strain level, which cannot be grappled with by intersecting higher-level taxon IDs with functional data (Douglas et al., 2020; Louca et al., 2018). GeoChip (He et al., 2007) has already been mentioned, but otherwise direct inference of function from sequences has primarily been performed from metagenomic data, although this is also amenable to target capture (Moore et al., 2018). Unfortunately, the acquisition of functional DNA directly from amplicon sequencing is largely infeasible as universal primers are generally impossible, even for conserved function (e.g., denitrification: Priemé et al., 2002; nitrogenase: Angel et al., 2018), creating a pseudoabsence problem.

As an example of an arena in which to compare methods, plant-microbe symbioses are particularly complex. Microbial communities in the rhizosphere, root, and root nodules are important to nutrient acquisition, plant growth, and confrontation of biotic and abiotic stress (Han et al., 2020). These communities consist of diverse bacteria and fungi that interact with the host plant and with each other to perform nitrogen fixation, hormone production, nutrient cycling, and pathogen suppression among other processes (Liu-Xu et al., 2024). The rhizosphere is highly diverse and generally dominated by the bacterial phyla Proteobacteria, Actinobacteria, Bacteroidetes, Firmicutes, and Acidobacteria, as well as the fungal phyla Ascomycota and Basidisomycota (Pantigoso et al., 2022). In the root, endophytic communities are generally smaller selective group of phyla deriving from the rhizosphere), usually dominated by Proteobacteria, Actinobacteria, and Bacteroidia (Ramula et al., 2024), while endophytic fungi are mainly root-associated Ascomycota, Basidisomycota, and Glomeromycota (Taniguchi et al., 2023). Root nodules, symbiotic organs that host nitrogen-fixing bacteria, are found in many plants and are important as highly specialized plant-microbial symbiosis organs. The microbiome environment of nodules is surprisingly poorly understood; these specialized organs are not just microbial landing zones for rhizobia and other nitrogen-fixing bacteria, but dynamic hubs of interaction where non-rhizobial nodule-associated bacteria (NAB) can act as co-pilots that help to enhance symbiotic efficiency and promote plant growth (da Silva et al., 2023). The root nodules of leguminous plants (family Fabaceae, including important crops like soybean (*Glycine max*) and alfalfa (*Medicago sativa*), as well as the non-leguminous *Parasponia* (syn. *Trema*) in Cannabaceae) are inhabited by both symbiotic nitrogen-fixing rhizobia (e.g., members of the genera *Rhizobium*, *Bradyrhizobium*, and *Ensifer* (syn. *Sinorhizobium)*) and non-rhizobial Proteobacteria (e.g., *Bacillus*) and Actinobacteria (Ali et al., 2024), while actinorhizal plants (from 8 other plant families) are inhabited by both *Frankia* and non-*Frankia* (Ghodhbane-Gtari et al., 2021). Fungal communities within root nodules are dominated by Ascomycota, with less abundance of Basidiomycota and Glomeromycota which indicate the strong host and niche filtering (Taniguchi et al., 2023). Nodule-colonizing fungi, such as *Aspergillus, Glomus, Penicillium, Trichoderma, Macrophomina, Fusarium,* and *Rhizoctonia*, have been isolated from legume root nodules using culture techniques (Adan et al., 2024). Overall microbial diversity decreases from rhizosphere to root to nodule, while Proteobacteria, Actinobacteria, and fungal Ascomycota to a lesser extent Basidiomycota phyla are present across rhizosphere, root and nodule (Ramula et al., 2024; Taniguchi et al., 2023), reflecting both common environmental reservoirs and strong host-driven selection.

The diversity and abundance of these communities are essential for studying symbiotic efficiency, host specificity, and composition of rhizosphere, root, and root nodule microbiome, yet such complexity yields methodological challenges. In particular, while inference of nitrogenase function commonly uses genus-level taxonomic designations (Boyd & Peters, 2013; Dos Santos et al., 2012), this methodology is far from perfect as nitrogenase function actually differs even at the strain level between particular species. The desire to study both fungal and bacterial members of the symbiosis, as well as their function, points to the need for a method more amenable to efficiently capturing multiplexed target loci, affording access to eukaryotic and prokaryotic community information with sequenced-based potential function information.

In this study, we have developed a multidomain triplet target hybridization capture method designed to enrich eukaryotic ribosomal markers (the ITS1, 5.8S, and ITS2 spacer region) using a novel probe panel, implemented as a multiplex experiment with the previously designed 16S panel, and to design a new functional panel representing core bacterial symbiotic genes (e.g., *nifH* and other nitrogenase subunits, as well as *nodA* and other non-nitrogenase genes that participate in other aspects of symbiosis). We designed and tested the performance of the overall capture kit in rhizospheric soil samples, roots, and root nodule microbiomes. This is to our knowledge the first study to design ITS probes for both plant host and fungal symbionts, as well as the first study to multiplex functional and taxonomic data acquisition from microbial communities in the form of a probe-based target capture kit. As well as the bioinformatic design, we prepared libraries and performed capture on a range of empirical samples from multiple plant host families to assess performance. Resultant capture libraries were compared with standard 16S rRNA and ITS amplicon libraries, both for the empirical samples and a mock community reference of known composition.

Specifically, our goal was to assess the experimental efficiency of novel ITS and functional gene probes, and to evaluate whether hybridization capture improves (i) detection of rare and low-abundance taxa, (ii) recovery of function symbiotic genes, (iii) detection of the fungal symbionts/endosymbionts, and (iv) accuracy of community structure estimation. By integrating this multidomain target enrichment technique, comprising bacteria, fungi, and plants, into a single workflow, this approach aims to provide a scalable, cost-effective method to study the microbial communities in the rhizosphere, root, and root nodules.

## METHODS

### Samples collection

#### Experimental success comparison

Samples were collected from a total of 36 distinct field sites from the NEON network of ecological observatories (https://www.neonscience.org/). Sample outcomes for the purpose of reporting (*n* = 2,971) were recorded as follows: DNA extraction yield, sequencing yield, and on-target yield for 16S, ITS, and symbiosis genes. For direct sequence analysis, including DNA assembly and microbial community analysis, two data subsets were taken and sequencing results are reported here in full (*n =* 44), the first testing microbial community estimation and the second testing the acquisition of functional genes.

#### Microbial community comparison

A total of 7 root nodule samples were collected from the following leguminous host plants: *Aeschynomene indica, Apios americana, Desmodium sessilifolium, Lackeya multiflora, Sesbania herbacea* (two replicates), and *Trifolium lappaceum* from local sites in Mississippi. These samples were chosen as the subject of a previous amplicon sequencing study (Pantinople et al., 2026) which affords a direct comparison with target capture. The species were also selected to represent a phylogenetically broad set of leguminous hosts that are typical eastern North American flora and involved in root nodule symbiosis.

### DNA Extraction

DNA extraction follows the protocol of Pantinople et al. (2026) and is outlined briefly here. All DNA extractions were performed under sterile conditions inside a PCR hood. Each sample was separated into rhizospheric soil, root, and root nodules, described as follows. Root samples were placed in a 2 ml microcentrifuge tube and molecular grade water was added with gentle tapping to dislodge adhering soil. The soil wash was centrifuged at 10,000 g for 5 min to pellet the rhizospheric soil. The root along with the root nodule was surface sterilized to eliminate epiphytic microorganisms by successive 1 min washes in 70% ethanol and 5% bleach (Johnson, 2019), and then rinsed twice with molecular grade water to remove residual bleach. Root nodules were dissected from root and transferred to 2 ml microcentrifuge tube while the remaining root tissue were transferred to another 2 ml microcentrifuge tube. Thus, one sample is converted into three samples for DNA extraction: (1) rhizosphere (“Rh”), (2) root (“Ro”), and (3) nodule (“No”). As a positive control for estimating diversity and abundance, mock communities were also submitted for all downstream analysis, with two replicates each for amplicon sequencing (see below) and target capture, using the Zymo Biomics DNA community standard (Zymo Research D6306, Irvine, CA).

DNA was extracted from rhizosphere, root and nodule samples using a column-based open source protocol (Pantinople et al., 2026; Williamson et al., 2014). Five hundred µL lysis buffer was added in the 2 ml microcentrifuge containing rhizospheric soil, root and root nodule and samples were mechanically disrupted with 4-6 sterilized metal beads in a tissue disruptor (FisherBrand Bead Mill 24; speed = 3.10 m s ¹, 4 cycles of 30 s with 5 s pauses) to lyse the cells. Samples were incubated at 65°C for 20min in a hot water bath, processed a second time in the tissue disrupter to ensure complete cell lysis, incubated a second time at 65°C for 10min, and centrifuged at 10,000 g for 2min. The supernatant was transferred to a new microcentrifuge tube, 200 µL potassium acetate was added to precipitate proteins and detergents. The samples were then incubated at -20 °C for at least 1 hour, centrifuged at 10,000g for 30 min, and the clarified supernatant was transferred to fresh tubes and combined with 1.2 mL guanidine hydrochloride. The mixture was loaded onto a silica spin column (EconoSpin®, Epoch Life Science, Missouri City, TX, USA) in 700 µL aliquots and centrifuged at 10,000g for 2 min, discarding the flow-through after each spin.

Columns were washed twice with 500 µL wash solution (10 mL 1 M Tris, pH 8.0; 2 mL 0.5 M EDTA; 10 mL 0.5 M NaCl; 670 mL ethanol; 308 mL molecular-grade water) and once with 80% ethanol, centrifuged at 10,000 g for 2 min after each wash. Columns were then centrifuged at 10,000 g for 5 min to remove residual ethanol and eluted in storage buffer (5 mL 1 M Tris, pH 8.0 in 495 mL molecular-grade water) after room temperature incubation for 10 min, followed by centrifugation at 10,000 g for 2 min. DNA concentrations were quantified using a Qubit fluorometer (Thermo Fisher Scientific, Waltham, MA, USA).

### ITS Panel Design for Target Capture

First, we replicated the clustering procedure of Beaudry et al. (2021), performing a USEARCH analysis on the entire UNITE fungi database (version 25.07.2023; Nilsson et al., 2019), finding that this yielded an excessive and uneconomical capture space (83,934 sequences with a capture space of 5,389,8499 bp, which would require hundreds of thousands of probes). Therefore we adopted a different approach from (Beaudry et al., 2021), observing that for most fungi the ITS1 and ITS2 pieces are individually very short, often <200 bp, which is less than a standard Illumina library insert size. These two regions are flanked by the 18S, 5.8S, and 28S rDNA genes, which are strongly conserved. We aimed instead to design a probe set that captures these conserved regions, so the probes were designed to match a much less variable flanking target with the aim of recovering the more variable ITS regions via the “splash zone” effect (Straub et al., 2012): the tendency of library fragments that are not targeted to be sequenced due to being within a distance of the standard insert size from the target and experiencing effectively *in vitro* sequencing linkage. This strategy closely resembles that used for many years to target variable non-coding regions surrounding captured UCEs (ultraconserved elements; Faircloth et al., 2012) and introns associated with exonic capture (Johnson et al., 2016).

To design the conserved target, we first conducted a USEARCH cluster (per Beaudry et al., 2021) of the entire UNITE fungi database with 0.9 identity and 0.9 coverage parameters with the aim to initially reduce the number of sequences. This yielded 83,934 sequences from 206,494 total, and the result was annotated using three GenBank ITS sequences (one Ascomycete, one Basidiomycete, and one Glomeromycota) and the Geneious Prime v. 2025.1.3 (Kearse et al., 2012) gene annotation tool at 70% identity (needed to accommodate variability in the large subunit). Annotated fragments were limited to the entirety of 5.8S (usually ∼160 bp), and flanking fragments of the large and small subunit, also limited to ∼160 bp to match the length of 5.8S annotations. Because UNITE sequences differ in length and gene coverage (Nilsson et al., 2019), this yielded 82,516 5.8S, 7,263 large subunit, and 3,551 small subunit sequences. Regions extracted by this annotation result were then clustered again at 0.9 identity and 0.9 coverage in USEARCH, yielding 4680, 561, and 302 clusters respectively, of which representative centroid sequences chosen by USEARCH were taken as the final capture space for probe design (cumulative probe length 884,241 bp; effective capture space estimated as aligned length: 2291 bp).

### Functional Panel Design for Target Capture

A database of functional nitrogen-fixing annotations was developed from a literature review we conducted, using the gene list reported in Table 4.1 in Thomas and Rahman (2020), updated with a series of reports of lineage specific genes from papers reported in Supplemental Table 2. The database includes *nif*, a functional gene cluster that includes several families of nitrogenase subunits, regulatory genes, and other genes relevant to nitrogenase activity; *nod*, a functional gene cluster including several families of genes required to nodulate plants, and further named accessory genes such as *noe* and *nol* (a full list of targets is included in Supplemental Table 2). This database was used to annotate regions from a set of genomic references to design the bait panel. Reference genomes were selected from GenBank on the basis of having a provenance deriving from a nodule and being a high-quality annotated chromosome-scale assembly. From taxonomic duplicates, one representative was chosen from each rhizobial genus, and several isolates of *Frankia* to represent the *Frankia* clusters (clades) that inhabit nodules. Selected were 20 genomes from 14 distinct taxa (Supplemental Table 1), including 9 alpharhizobia, 2 betarhizobia, and 3 *Frankia.* Selected assemblies were downloaded in Geneious Prime 2025.1.3 and searches were conducted on the reference gene name list including synonyms recognized by GenBank. Regions containing these genes were annotated manually as “symbiosis islands,” which were defined as clusters of nodulation-relevant genes in contiguous clusters, allowing inclusion of other annotated genes if they were no more than 1000 bp apart from and between target genes (beyond this criterion islands were annotated separately). Both plasmids and the main bacterial chromosome were searched for matches. In a small number of cases, symbiosis-relevant genes were isolated from these clusters with no neighboring relevant genes and were individually annotated. This resulted in 85 distinct regions from the 14 taxa (mean length 3,958 bp, minimum 111 bp and maximum 34,967 bp) representing 73 distinct genes (18 *nif* genes, 17 *nod* genes, and the remainder representing accessory genes such as *fdx, fix, noe,* and *nol* genes) with a cumulative probe length of 336,388 bp and effective capture space estimated by alignable length as 323,447 bp.

### Probe design

Probe design was conducted by Daicel Arbor Biosciences (Ann Arbor, MI, USA) by tiling 80 bp probes at 3× coverage of the target regions. The probes were then screened for repeat regions that were then masked. This led to 38,406 probes of which 59.5% covered ITS and 40.5% covered the functional loci.

### Library Preparation and Target Enrichment

Libraries were prepared using the NEBNext Ultra II kit (New England Biolabs). Based on the nature of eDNAs from very small samples (nodules < 1 mm are common and multiple structures were not generally combined), initial optimizations, and observed sample quantification and size distribution, input DNAs were divided into two processing paths. (1) Extractions that exceeded 250 ng total were normalized to 500 ng (or the entire sample was volume-normalized if this could not be achieved). These samples were then sonicated (8 cycles of 30 s medium power sonication, 90 s rest on a Bioruptor machine (Diagenode, Denville, NJ, USA)) and libraries were prepared in miniaturized half reactions. (2) Extractions with 250 ng or less were not sonicated (it was observed that, for these samples, degradation and yield were closely correlated so that quantification could serve as a size distribution proxy), and the samples were subjected to miniaturized quarter reactions (both to conserve resources and to normalize adapter input to sample quantity). Miniaturization was applied at the end prep and adapter ligation and cutting phases, with all downstream steps (bead washes and PCR enrichment) brought to half volume as in path (1) for sample handling purposes. Samples in path (1) were subject to 13 PCR cycles while samples in path (2) were subject to 15 cycles.

Samples were quantified and pooled on an equimolar basis to yield 24- to 28-plex pools (aiming for 864 to 1008 ng total DNA), and sample pools were structured based on library yield to group underperforming libraries and prevent undesirable *in vitro* competition. Generally, all libraries that had less than the desired input (32 ng plus pipetting error adjustments) were grouped, and in a few cases near-32-ng and lower samples were separately grouped. Probe-based sequence capture was conducted according to the v5 MyBaits protocol with these exceptions: the 16S and ITS/functional probe sets were pooled on an equivolume basis for multiplexed hybridization reactions, the hybridization temperature was 65°C, and the hybridization length was 36 hr. The final post-capture PCR enrichment was also divided among sample workflows, with 13 cycles for pathway (1) and 15 cycles for pathway (2). Finally, post-capture pools were cleaned with 0.8× SPRI beads, quantified, and pooled again on an equimolar basis to yield 384-plex pools for sequencing. All pools were sequenced by Novogene (San Diego, CA) on a NovaSeq X Plus 10B lane (paired end 150 bp reads).

### Community diversity comparisons

Amplicon sequencing results were prepared from a set of 21 samples also prepared for target capture as described above. These are previously reported data (Pantinople et al., 2026) and are described briefly here. Bacterial 16S v4 amplicons were prepared using primers 515f and 806r (Thompson et al., 2017) and fungal ITS1 amplicons were prepared using ITS1-F (Gardes & Bruns, 1993) and ITS2 (White et al., 1990) with full reaction conditions reported in Pantinople et al. (2026). Individual libraries were prepared from these and sequenced on a MiSeq instrument at the Michigan State Research Technology Support Facility (East Lansing, Michigan, USA). Extensive attempts were made to also prepare comparable functional locus primers using both custom degenerate designs we generated and previously reported primer sequences (Tu et al., 2016; Zhao et al., 2020) and to optimize reaction conditions, but reliable amplification across samples was impossible due to a lack of conserved sites in both *nodA* and *nifH*. In general, application of amplicon sequencing to rapidly evolving functional loci is infeasible for taxonomically diverse samples. Thus, sequencing methodology comparisons focus on taxonomic identification in ITS and 16S.

### Community data analysis

Raw target capture DNA sequences were first sorted to individual capture sets using BWA (Li & Durbin, 2009). These sorted datasets were used both for downstream analysis and estimates of on-target percent. Based on optimizations, sequences were used directly for Kraken2 (Wood et al., 2019) analyses (below) and trimmed for adapter content and low-quality base calls using Trimmomatic v. 0.33 (Bolger et al., 2014) for SPAdes assembly (also below).

Sequence capture analysis used Kraken2 (Wood et al., 2019) with the SILVA database (v. 138) for full-length 16S (Chuvochina et al., 2025), and the UNITE database version for all eukaryotes (v. 19.02.2025) to analyze the ITS and flanking rDNA data. Kraken output was summarized with Bracken (Lu et al., 2022) and imported into R for analysis in the *phyloseq* package (McMurdie & Holmes, 2013). Amplicon taxonomic analysis followed previously reported methodologies (Pantinople et al., 2026), briefly including analysis in QIIME2 using DADA2 (Callahan et al., 2016) and a naïve Bayes classifier based on GreenGenes (DeSantis et al., 2006; McDonald et al., 2012). For both analytical pipelines, host DNAs, which were a minority of the data, were eliminated bioinformatically using mitochondrial and chloroplast classifications appropriate for either taxonomic database.

Analysis of mock community data used both correct identification and correct estimates of prevalence reported by the manufacturer as the relevant benchmarks (note these are not barcoding DNA dosage benchmarks but gDNA benchmarks, and therefore also represent DNA prevalence assumptions in abundance data). Analysis of real communities compared taxonomic content and abundance.

### Functional data analysis

Functional gene data from the symbiosis island capture kit were analyzed on a subset of 21 samples. These were chosen among the NEON samples rather than the amplicon reference, as the amplicon comparison was not possible and a wider range of hosts was needed to contrast gene content. Only nodule samples were chosen as the detection of rare functional gene sequences from soil samples was not of specific experimental interest (see Results). The target host taxa included the legume genera *Aeschynomene, Alysicarpus, Amphicarpaea, Astragalus, Baptisia, Chamaecrista, Clitoria, Crotalaria, Dalea, Indigofera, Lespedeza, Macroptilium, Melilotus, Mimosa, Pediomelum, Pictetia, Tephrosia, Thermopsis, Trifolium,* and *Zornia.* We assembled the on-target functional sequences and shotgun sequences using SPAdes (v. 3.14.0) with default settings. Based on optimizations, the -meta SPAdes option (Nurk et al., 2017) produced the best output in terms of contig length and was used for all downstream analysis. Output was then directly processed to predict open reading frames (ORFs) using Prodigal (v.2.6.3) (Hyatt et al., 2010). The custom database was constructed for all targeted symbiotic genes (Fig.7) from the publicly available NCBI protein database. The predicted protein sequences were compared against a custom protein database and similarity searches were performed using BLASTP (v.2.16.0). For each gene within each sample the best hit was selected based on percent identity ≥ 90%; when no hit met the identity threshold the first best hit was retained to avoid false negative results. Filtered results were compiled across the 21 samples to generate a gene presence-absence matrix, and a heatmap was generated and visualized using pandas (v.3.0.0) (McKinney, 2011), matplotlib (v.3.10.8) (S. Han & Kwak, 2023), and seaborn (v.0.13.2) (Waskom, 2021).

### Data availability

The probe design, together with code to reproduce these analyses and figures, together with metadata and machine-readable community data files, are available at GitHub (https://github.com/ryanafolk/target_capture_paper). Raw sequence reads have been deposited at the Sequence Read Archive (Bioproject PRNJXXXXXXX).

## RESULTS

### On target success

Across the 2,971 samples assessed for experimental success, the average sequencing effort was 8.6 million reads. The average overall experiment success was assessed as percent of sample matching the target loci; this was on average 92.2%. Among the three distinct locus panels, the average on-target percent was 55.8% for 16S, 21.30% for ITS, and 15.16% for the symbiosis genes (Fig. 2D, E, F). This result closely reflects the input probe dosage of 5:3:2. ITS and symbiosis genes were sensitive to background DNA frequency as they differed strongly by sample type (Fig. 2E, F); ITS recovery was maximal in root tissue (31.6% average on-target for all root samples) and symbiosis gene recovery was maximal in nodule samples (33.0% average on-target). Hybridization efficiency was also assessed across host taxa at the level of family; 16S and ITS were highly consistent across families (Fig. 2A, B), while symbiosis gene recovery was sensitive to host taxon (Fig. 2C), perhaps reflecting either divergent symbionts or differing gene content among symbionts.

**Fig. 1.**
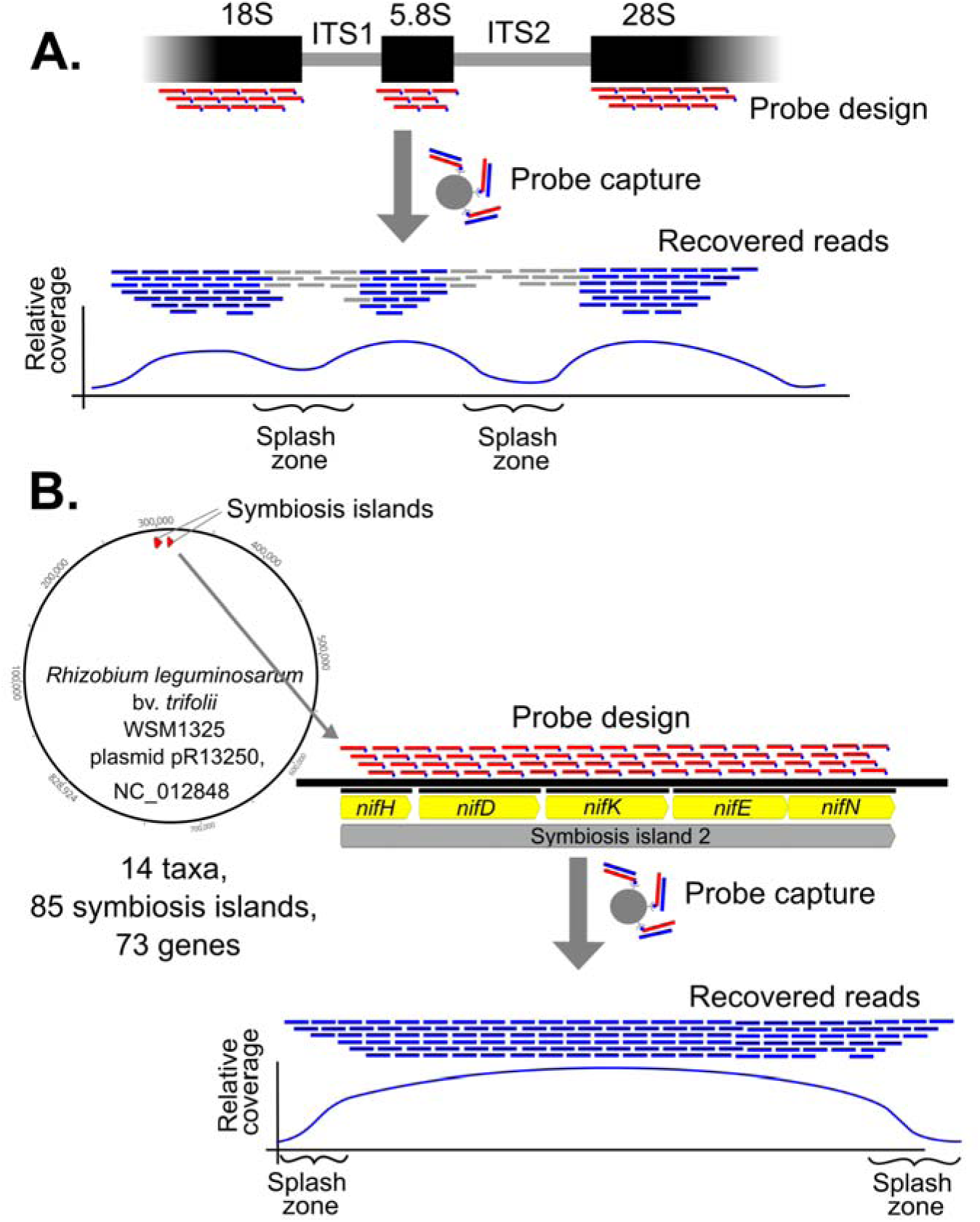
Conceptual design of ITS barcoding (A) and symbiosis gene (B) kits. Tiled synthesized probes are represented in red, recovered library molecules in blue (on-target) and gray (off-target), and gene models in yellow. Coverage graphs conceptually indicate the existence of “splash zones,” regions not targeted but recovered in the output by virtue of an insert size larger than the probe hybridization length.

**Fig. 2.**
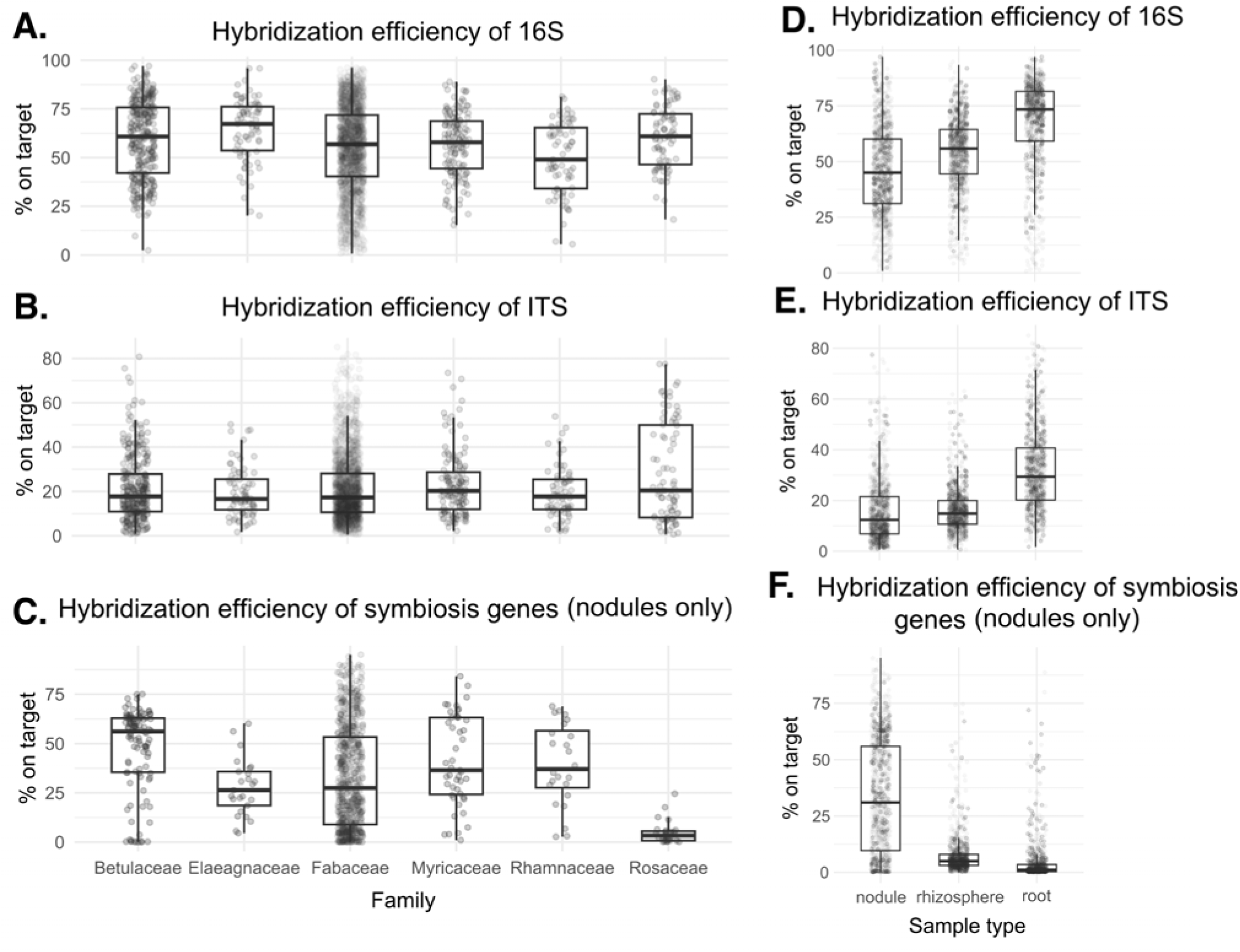
Hybridization efficiency for 16S (A, D), ITS (B, E), and symbiosis gene (C, F) targets across host families (A-C) and sample types (D-F). The symbiosis gene panel is sensitive to the background frequency of the target genes, but otherwise performance was similar across samples.

Because our experiment was designed as a highly multiplexed targeted experiment, going after three functionally and taxonomically distinct locus classes, assessing the relative performance of these was of interest. Regressions (Fig. 3) indicated that 16S and ITS were positively correlated with modest strength or relationship (Fig. 3A). Thus samples tend to perform similarly for both of these locus sets. By contrast, the symbiosis genes tend to perform inversely to 16S (not shown; see GitHub for input data) and ITS (Fig. 3B). Because symbiosis gene performance was sensitive to taxon and sample type (Fig. 2), this relationship is likely confounded by background frequency heterogeneity (e.g., functional symbiosis genes are probably present but very rare in rhizospheric soil compared to background genomic DNA), but because success in a multiplexed sequencing experiment is zero-sum this shows up as a negative correlation and not necessarily evidence of *in vitro* conflict between the assays. The average on-target percent data indicate >1 million reads per locus set for the typical sample, well above community standards for barcoding eDNA samples.

**Fig. 3.**
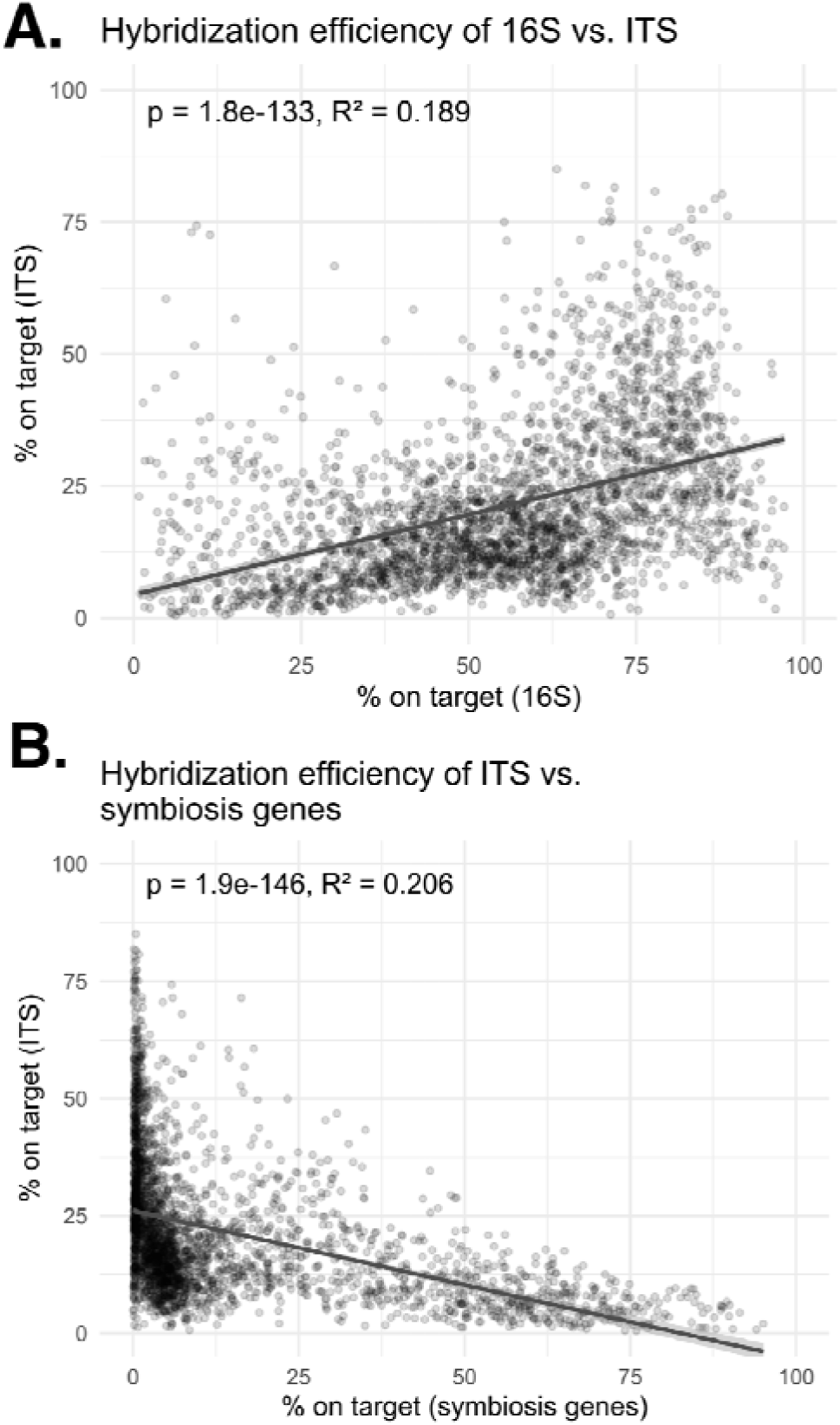
Comparative performance of multiplexed target capture. (A) The two DNA barcoding kits (16S and ITS) showed a positive, moderate correlation. (B) Barcoding kits (ITS shown here, 16S similar) were moderately negatively correlated. Given the strong dependence of the symbiosis gene kit on sample type (Fig. 1F), these results likely indicate the symbiosis gene panel is sensitive to background frequency rather than in direct in vitro conflict with the barcoding kits.

We compared microbial community composition across plant host tribes and sample types (rhizosphere, root, nodule) using amplicon sequencing and multidomain triple target capture (Figs. 4-5). Across both bacteria (16S) and fungi (ITS), the community composition variation was primarily methodological.

**Fig. 4.**
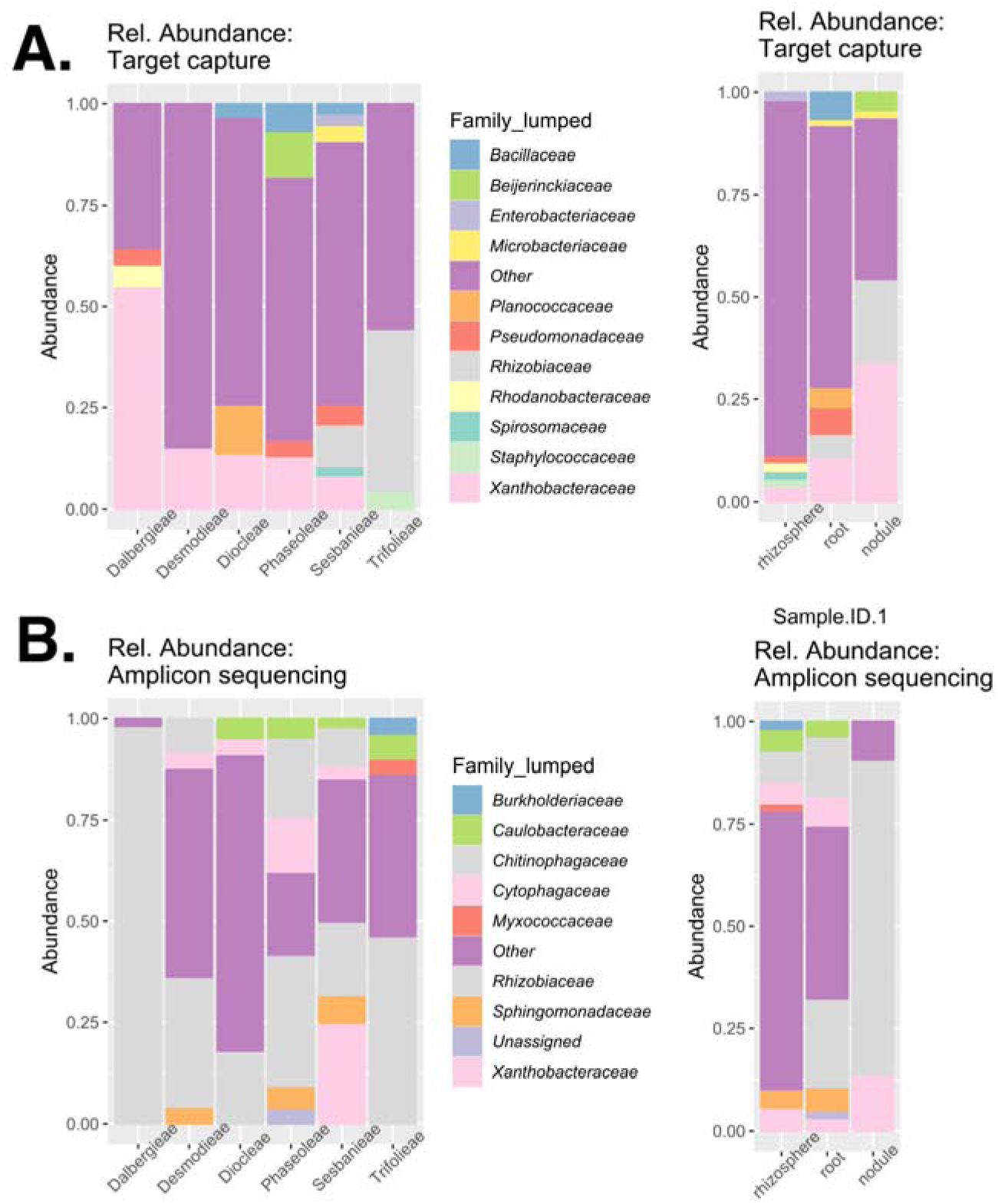
Relative abundance of bacterial families (16S calls) for (a) target capture and (b) amplicon sequencing, aggregated to plant host taxon (left column) and sample type (right column). Category “Other” compresses rare taxa (all taxon calls at the per-sample level that are < 0.5% relative abundance).

**Fig. 5.**
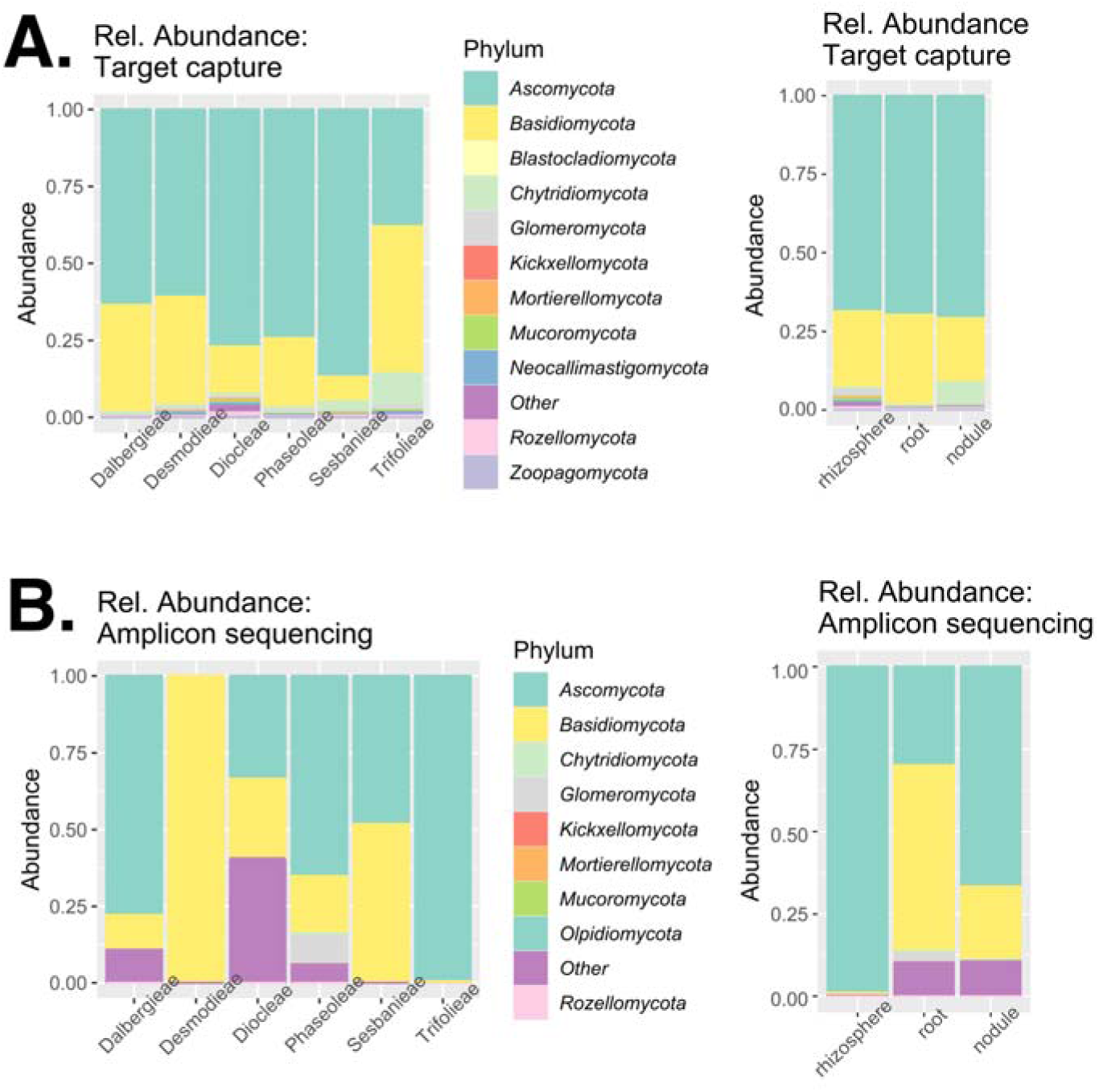
Relative abundance of fungal phyla (ITS calls) for (a) target capture and (b) amplicon sequencing, aggregated to plant host taxon (left column) and sample type (right column). Category “Other” compresses *incertae sedis* taxa and (mostly) undetermined amplicon sequence calls.

For bacteria (Fig. 4), both techniques consistently detected the dominant nitrogen-fixing families, particularly Rhizobiaceae and Xanthobacteraceae (Padda et al., 2022) but the relative abundance and detection of minor taxa differed significantly. Amplicon sequencing yielded a microbial community heavily dominated by a narrow set of families particularly in low density root and nodule samples where maximum percentage of reads were assignable to dominant taxa (Fig. 4b, right). In contrast, target capture reduced the dominance while simultaneously detected uncovered minor taxa, including Bacillaceae, Pseudomonadaceae, and Rhodanobacteraceae (Fig. 4a), reported in the root nodule-associated bacteria isolation studies (Pang et al., 2021). Uniquely, target capture recovered the nitrogen fixing Beijerinckiaceae (Borken et al., 2016; Padda et al., 2022) in the rhizosphere, which was absent in amplicon sequencing. These results indicate that the amplicon sequencing method may produce possible artifactual estimates of dominance in low diversity samples (Jia et al., 2022; Qin et al., 2023) target capture provides a more balanced representation of the community structure.

Similar methodological shifts were observed for the mycobiome (Fig. 5), where target capture consistently provided higher resolution compared to amplicon sequencing. Fungal communities were dominated by Ascomycota and Basidiomycota as a major dominant taxa (Ben-Laouane et al., 2026; Wang et al., 2024; Wen. Yang et al., 2023; Yang et al., 2024; Zhao et al., 2022) in amplicon sequencing while target capture recovered Ascomycota and Basidiomycota as most abundant but also detected Basidiomycota in the Trifolieae host tribe and the minor taxa Glomeromycota, Chytridiomycota, Mortierellomycota, and Rozellomycota across the samples (Xu et al., 2012) indicating enhanced sensitivity to rare taxa. Across both bacteria and fungi, target capture reduced the estimated dominance of most-abundant lineages and detected low-abundance and uncovered taxa across host tribes and sample types. These patterns suggest greater sensitivity of the multidomain target capture approach for detecting rare and uncovered microbial diversity.

### Community diversity comparisons by mock community resequencing

The mock community comprised ten expected genera, eight bacterial (*Bacillus, Enterococcus, Escherichia, Lactobacillus, Listeria, Pseudomonas, Salmonella,* and *Staphylococcus*) and two fungal (*Saccharomyces* and *Cryptococcus*). Amplicon sequencing recovered the major bacterial phyla present in the mock community *Firmicutes* and *Proteobacteria*, but did not detect the less abundant fungal phyla *Ascomycota* and *Basidiomycota* (which should be detectable via mitochondrial reads). Target capture recovered both bacterial and fungal phyla. At the genus level, amplicon sequencing recovered seven genera *Bacillus, Enterococcus, Escherichia, Lactobacillus, Listeria, Pseudomonas,* and *Staphylococcus* but failed to detect one abundant bacterial genus, *Salmonella*. Target capture recovered all ten expected genera, which includes both most abundant bacterial genera and the less abundant fungal genera *Saccharomyces* and *Cryptococcus.* No method perfectly matched input abundances, which could be affected by not only wet lab factors but differing barcode DNA dosage in the reference sample through 16S copy number variation and differing background genome size. However, the relative abundance profile produced by target capture somewhat more precisely reflected the expected composition of the mock community and captured both major and minor fungal and bacterial genera with no missing taxa.

### Capture and annotation of functional loci

The functional gene assemblies were successful for all 21 samples. *nifD* and *nifK*, the two nitrogenase subunits, were detectable without exception, indicating the functional capacity for nitrogenase activity. Most accessory *nif* genes were also detected in all samples; *nifM* was not detected in any sample and this accessory gene was not included in the original bait design (however *nodE* and *nodT*, both species-specific genes, were successfully detected despite not being in the capture set; *nifM* may often exist outside a canonical symbiosis island). Genes of the *nod* cluster were more variable, with a *nodE, nodL, nodS, nodT,* and *nodZ* homolog variably detectable. *nifH* and *nodA* are the most common targets in phylogenetic and molecular ecology studies; these were always detectable. Accessory genes outside this cluster were highly variable, with only an *fdxN, fixU, nfeD,* and *nolB* homologs detectable across all samples.

## DISCUSSION

### Cost efficiency

Our implementation of target capture was highly cost-effective. The price of target capture experiments is sensitive to project size, as it requires a minimal up-front investment for probe synthesis. At our project scale, inclusive of library preparation, capture, and sequencing we achieved a per-sample cost of $22.34 (2025 prices). This is only slightly more than the price of amplicon sequencing at market rates (our average costs are $20.78 per sample), and results in far more data. Our amplicon sequencing projects yield ∼200,000 reads on average (Pantinople et al., 2026), whereas 8.6 million reads was the average sequencing effort in this experiment. Thus the per-read price difference is 40-fold, leaving aside that amplicon sequencing is most commonly performed on a per-locus basis. We did not perform a direct economic comparison with unselected metagenomic techniques, but generally multiplexing as heavily as we did, 384-plex per lane, would lead to lower coverage of samples and a greater likelihood of missed community members. High sequencing depth is important for fully capturing community richness and rare features, with the need for high sequencing effort generally scaling with community complexity (Gweon et al., 2019).

### Overcoming sample challenges

Target capture is also a valuable technique because it naturally lends itself to degraded DNA material, as has long been appreciated for museum specimens (Bi et al., 2013; Folk et al., 2015). Degradation of nucleic acids is also a typical feature of eDNA (Brandão-Dias et al., 2025), which tends to confound amplicon sequencing as it induces breaks between priming sites. In accordance with this, our nodule samples from amplicon sequencing had an 84.8% success rate as measured by visible gel bands; success was as low as 43.7% for rhizospheric samples and 38.7% for non-nodular root materials, which have lower DNA dosage (Pantinople et al., 2026). In the target capture experiment, unilateral amplification failure is not typical and instead reads display sequencing effort variance. Considering 200,000 16S reads as a floor for sample success, 95.4% of samples (including more challenging root and rhizosphere samples) were successful. Thus target capture is a promising technique for microbial community characterization from challenging samples, and would likely be straightforward for historical samples, although that was not directly tested here. Similarly, non-microbial samples could be adapted in a straightforward way with this protocol. For instance, in a forthcoming contribution, we will describe our success with pollen metabarcoding using this target capture approach.

### Lower bias

One of the most concerning aspects of amplicon sequencing is the potential for amplification bias. This manifests in two ways: through primer mismatch such that primers fail to be universal (Rathod & Silverman, 2026), and through competitive effects where the abundance of some elements of the community are exaggerated through amplification bias (Polz & Cavanaugh, 1998). Beaudry et al. (2021) first suggested that target capture exhibits lesser bias. The mechanisms for this are generally two: (1) while amplification is also required for sequence capture, fewer cycles are generally implemented and this is done through adapter primers that are guaranteed to be universal for functional library constructs and which do not exhibit 5’ overhangs, degeneracy, or other primer features that reduce efficiency. Additionally, (2) sequence capture responds holistically to sequence divergence over the whole probe and is not sensitive to single mismatches; the specificity of the probes can modulated by the reaction temperature (but see Paijmans et al., 2016) and under a standard reaction condition 70% sequence similarity is tolerated. In accordance with the results of Beaudry et al. 2021), while both amplicon sequencing and target capture exhibit bias, abundance bias was often less for target capture and target capture never exhibited missing community members. Abundance bias appears to be particularly problematic for nodule samples. Rhizobia tend to be monodominant community members as this is a microbial housing structure (da Silva et al., 2023; Gage, 2004; Martínez-Hidalgo & Hirsch, 2017), but estimates of 95% and 99% relative nodule abundance as we customarily estimated from amplicon sequencing are unlikely to be accurate. The highest abundance we measured for potential diazotrophic bacteria using target capture was just over a majority (Fig. 4a), more likely to be accurate given the abundance of non-diazotrophic microbes in nodules (da Silva et al., 2023) as well as plant host DNA in nodular tissue (Adan et al., 2024).

### Functional interrogation

The final benefit of our target capture approach is one generally out of reach for amplicon sequencing experiments: directly sequencing functional loci. Short of performing direct metabolic experiments, sequencing of the full complement of functional genes is the most direct way to demonstrate the potential for microbial function (Denonfoux et al., 2013). For nodulating diazotrophic bacteria, that generally means demonstrating the presence of *nif* and *nod* genes, which respectively show the presence of functional nitrogenase activity, and the ability to communicate with the plant host and establish infection. Amplification of *nif* genes is customary in amplicon sequencing experiments (Gaby & Buckley, 2012), often to the exclusion of 16S or another locus that would give information on non-diazotrophic community elements. The nitrogenase subunits are themselves not suitable for universal primer design, so *nifH* (an electron donor that is part of the nitrogenase complex) is generally targeted (Angel et al., 2018; Gaby & Buckley, 2012). Amplification of *nod* genes is less typical as there is no one family universal to nodulating bacteria and none of the reported primers for common *nod* families performs well in diverse samples, nor do custom primers, as we noted in Methods. As noted above, sequence capture relaxes the need for highly conserved sites, and replicating representation of genes from selected taxa can overcome very high divergence, meaning that target capture lends itself to capturing and sequencing functional genes. Our results indicate two important things, that (1) large numbers of genes from disparate families may be easily captured from samples, often as long contigs representing entire operons or gene islands (Fig. 7), and that (2) this functional characterization may be multiplexed with a barcoding locus for higher-resolution community characterization and better sequencing costs.

**Fig. 6.**
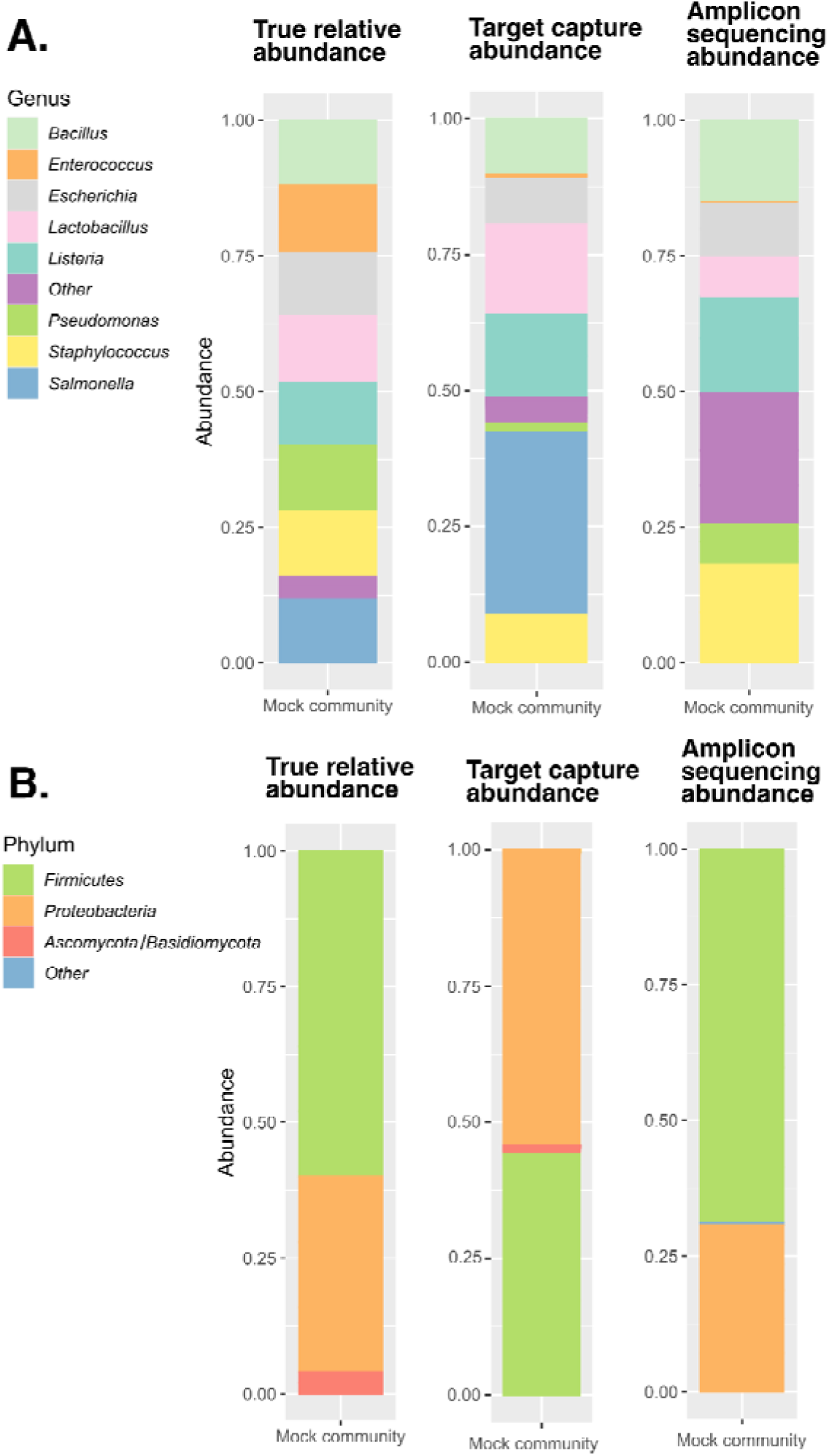
Comparison of mock community composition (true value in terms of total gDNA, left column) as estimated by target capture (middle column) and amplicon sequencing (right column). Both methods showed similar bias at the genus level (A) but only target capture detected all species in the mock community. At the phylum level (B) only target capture could detect both fungi and bacteria with 16S data.

**Fig. 7.**
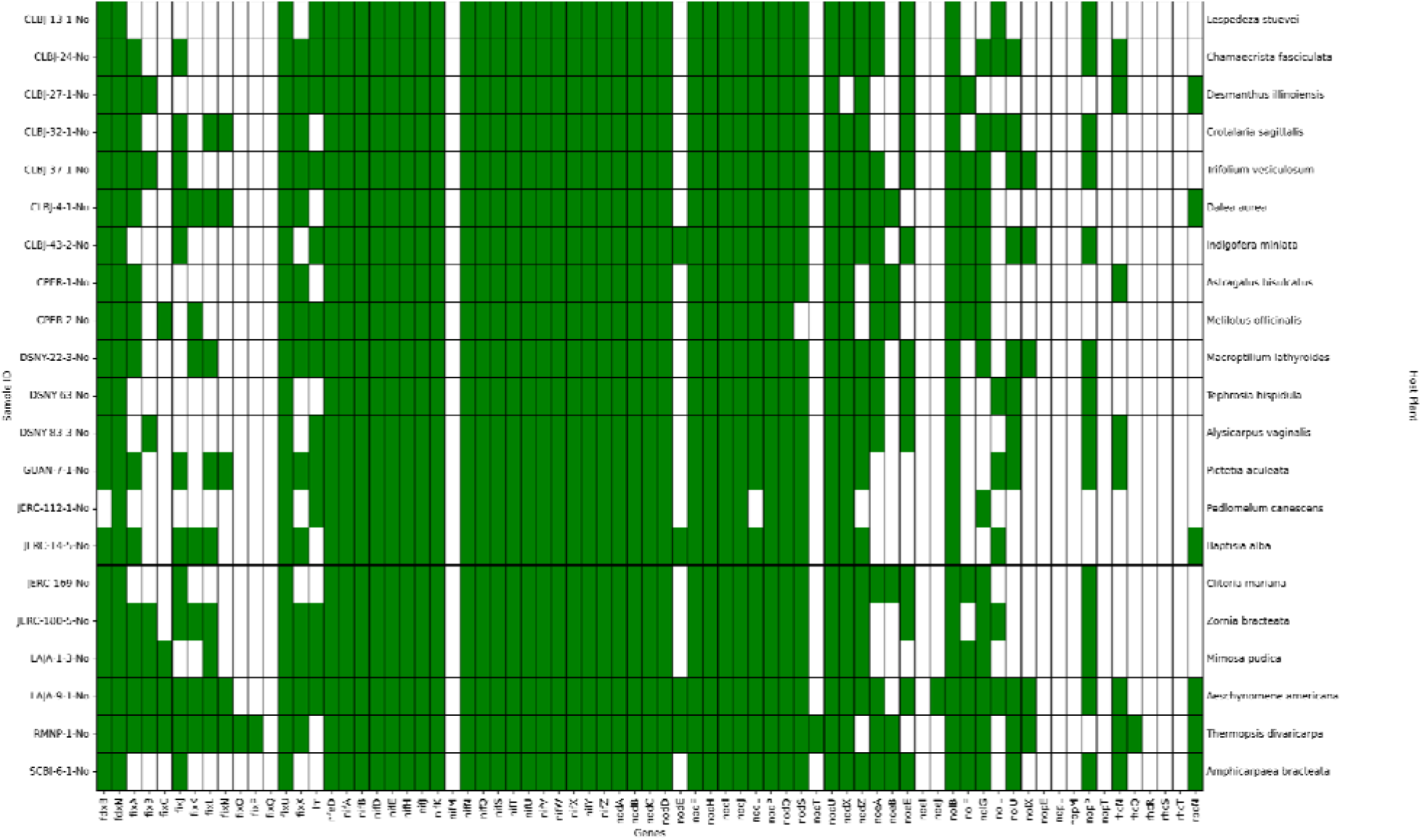
Heatmap illustrating the presence (green) and absence (white) of core nitrogen fixation genes targeted by functional gene probes. The x-axis represents a survey of named genes included in our survey; most are included in the kit, but compare (Supplemental Table 2).

### Conclusions

We designed a novel eukaryotic ITS barcoding kit for sequence capture as well as a kit for targeting loci on diazotrophic symbiosis islands. We multiplexed this with a previously reported 16S capture kit, evaluating capture success and community characterization success as compared to amplicon sequencing. We found that multiplexed sequence capture shows success over a range of sample types and qualities derived from plants and their environment. Sequence capture resulted in a less biased characterization of communities at a much stronger per-sample and per-read cost, as well as retrieving dozens of symbiosis-relevant genes in long, often contiguous symbiosis island contigs. Target capture is an important and underutilized tool in microbial ecology.

## Acknowledgments

Brian Brunelle and Megan Beaudry are thanked for discussions about the ITS probe design concept. This work was funded by the National Science Foundation, DEB-2316266.

---

**Supplemental Table 1.**
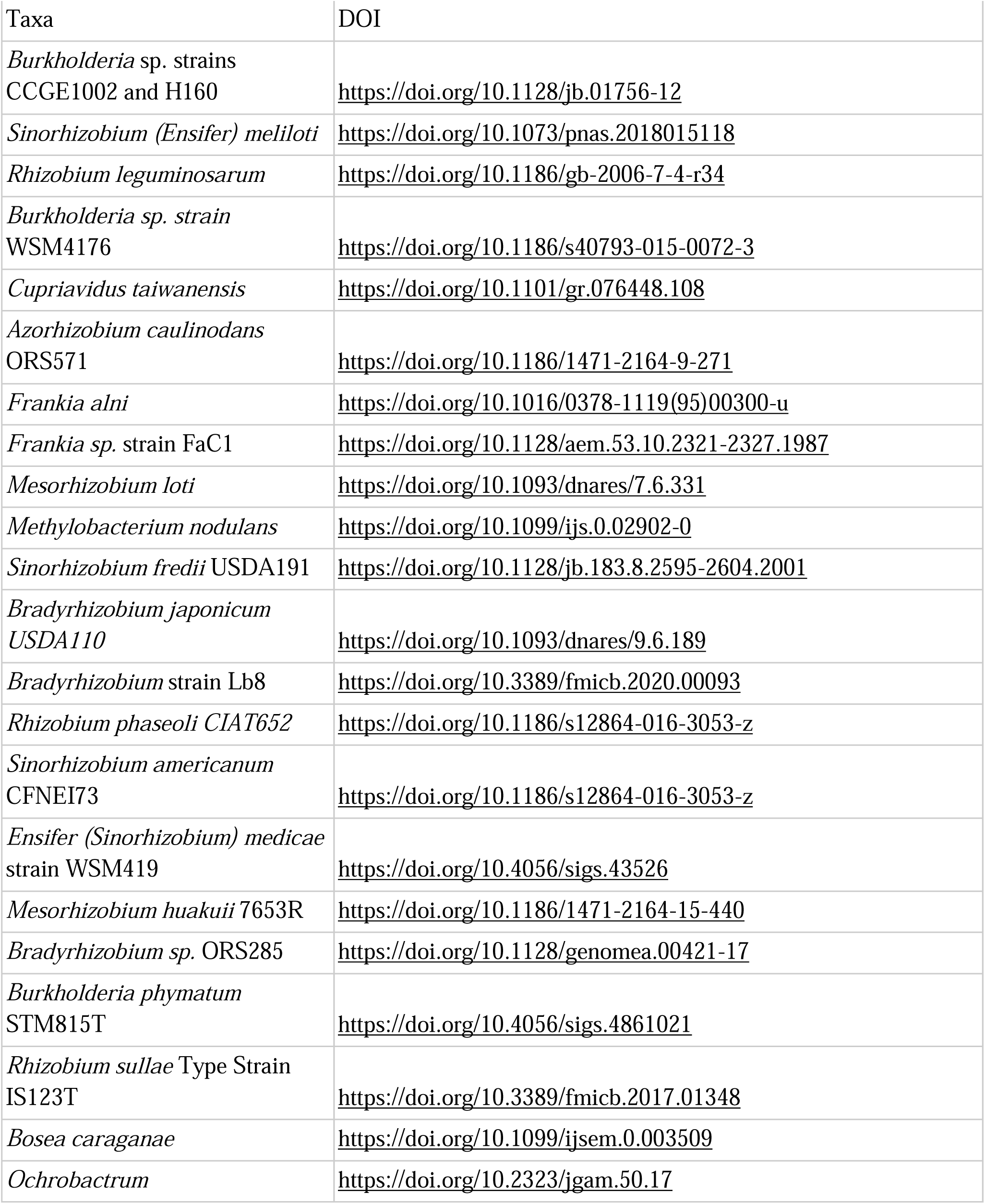
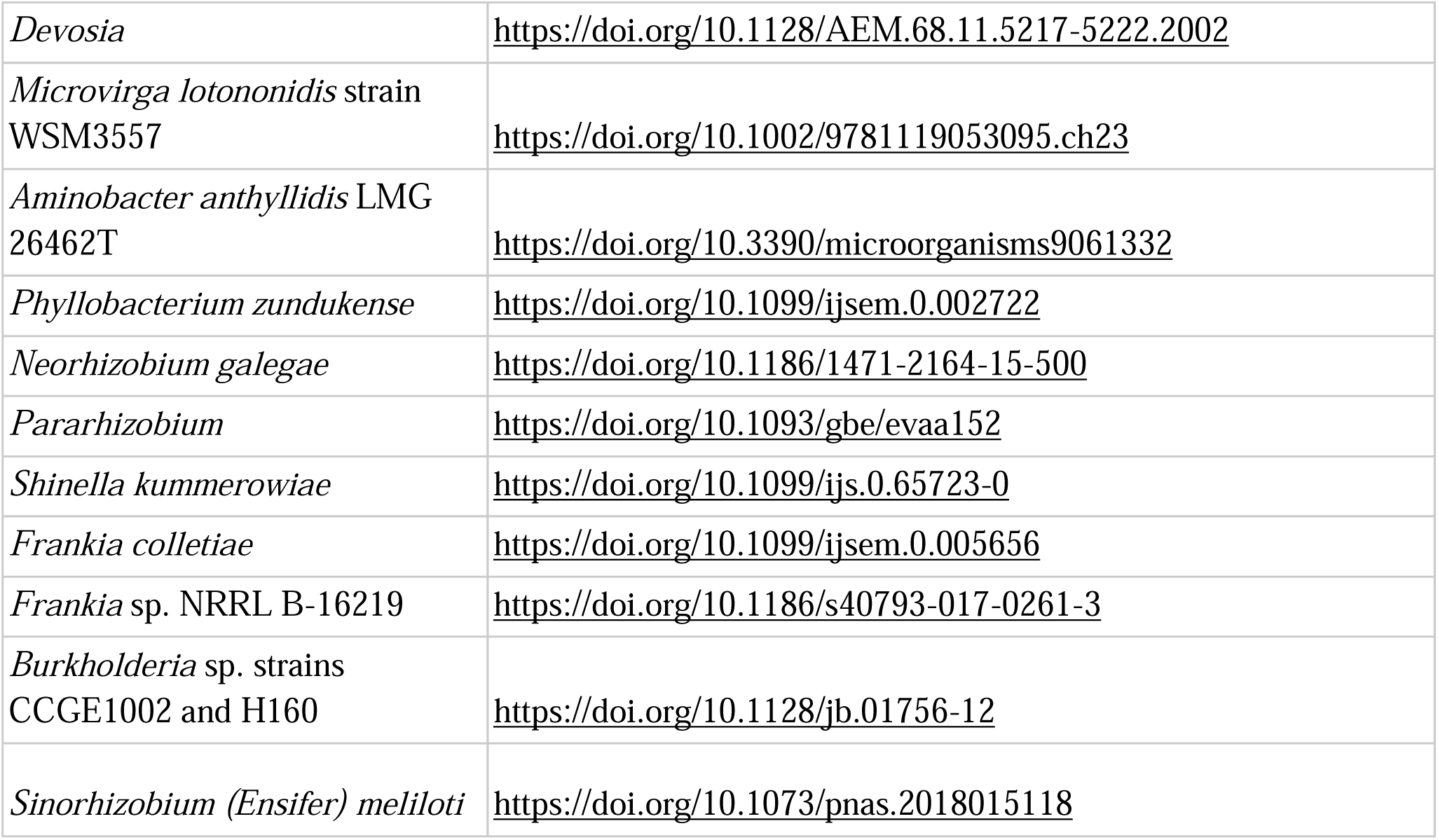
Literature review for retrieving gene annotations.

**Supplemental Table 2.**
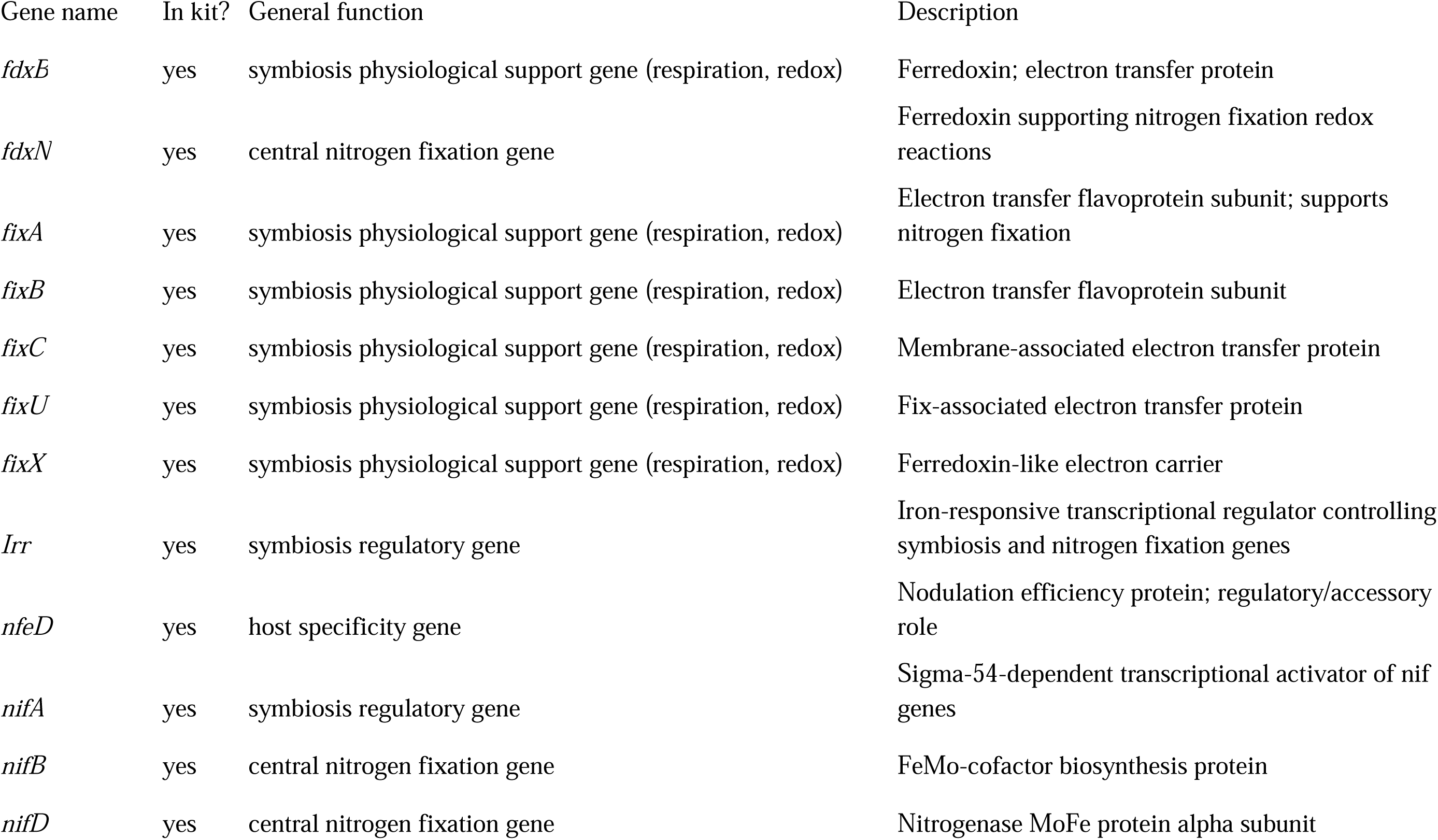
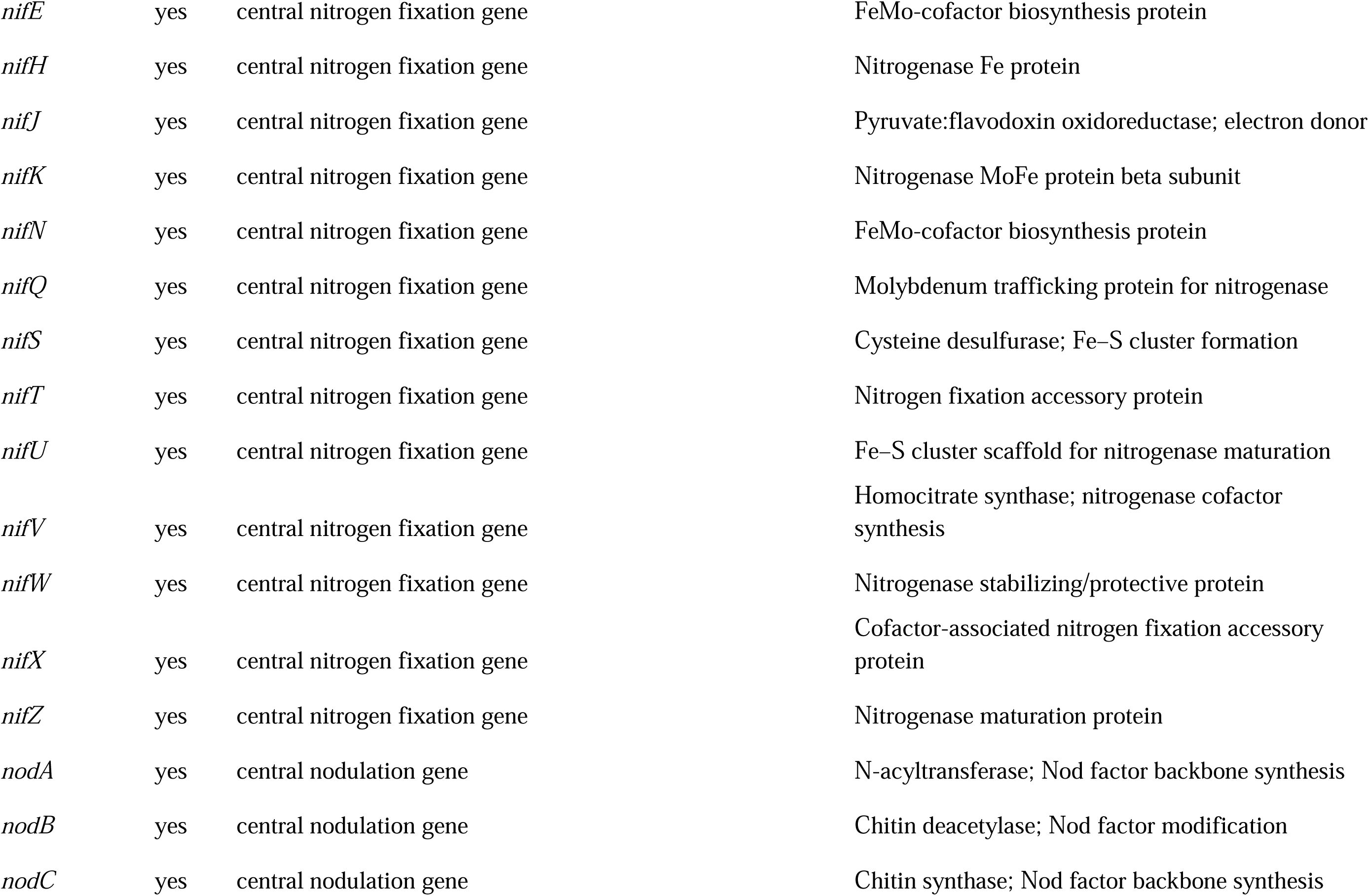
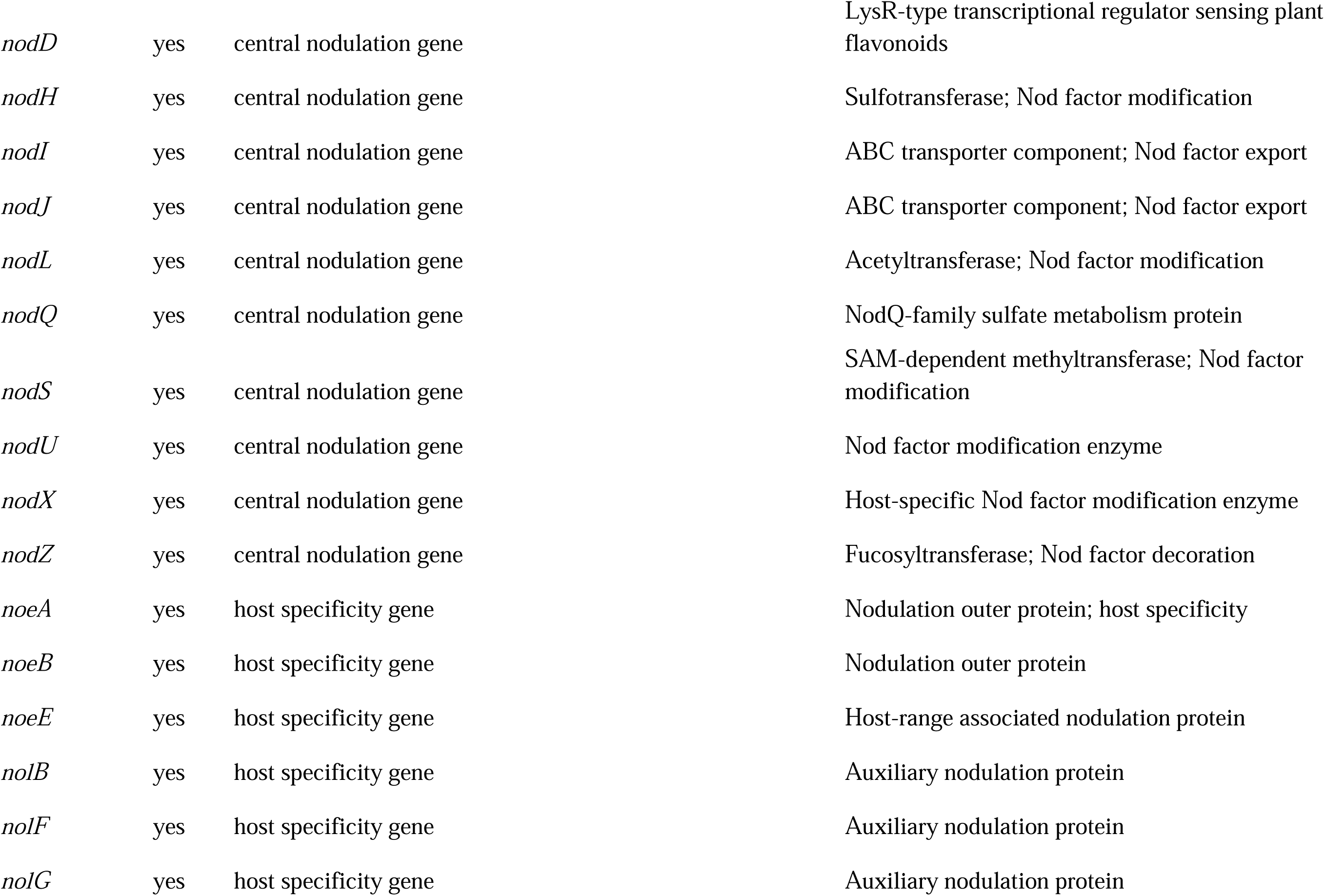
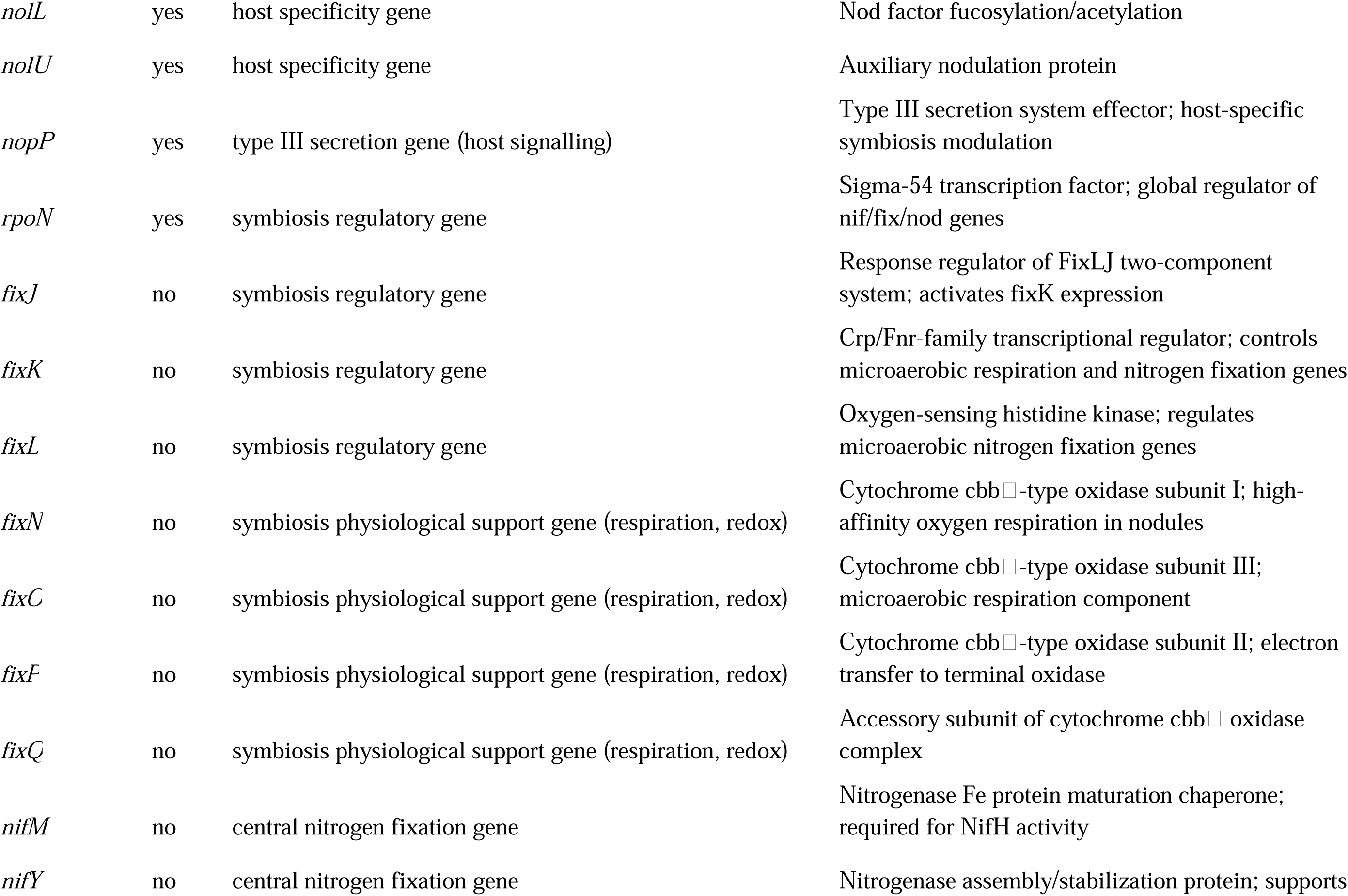
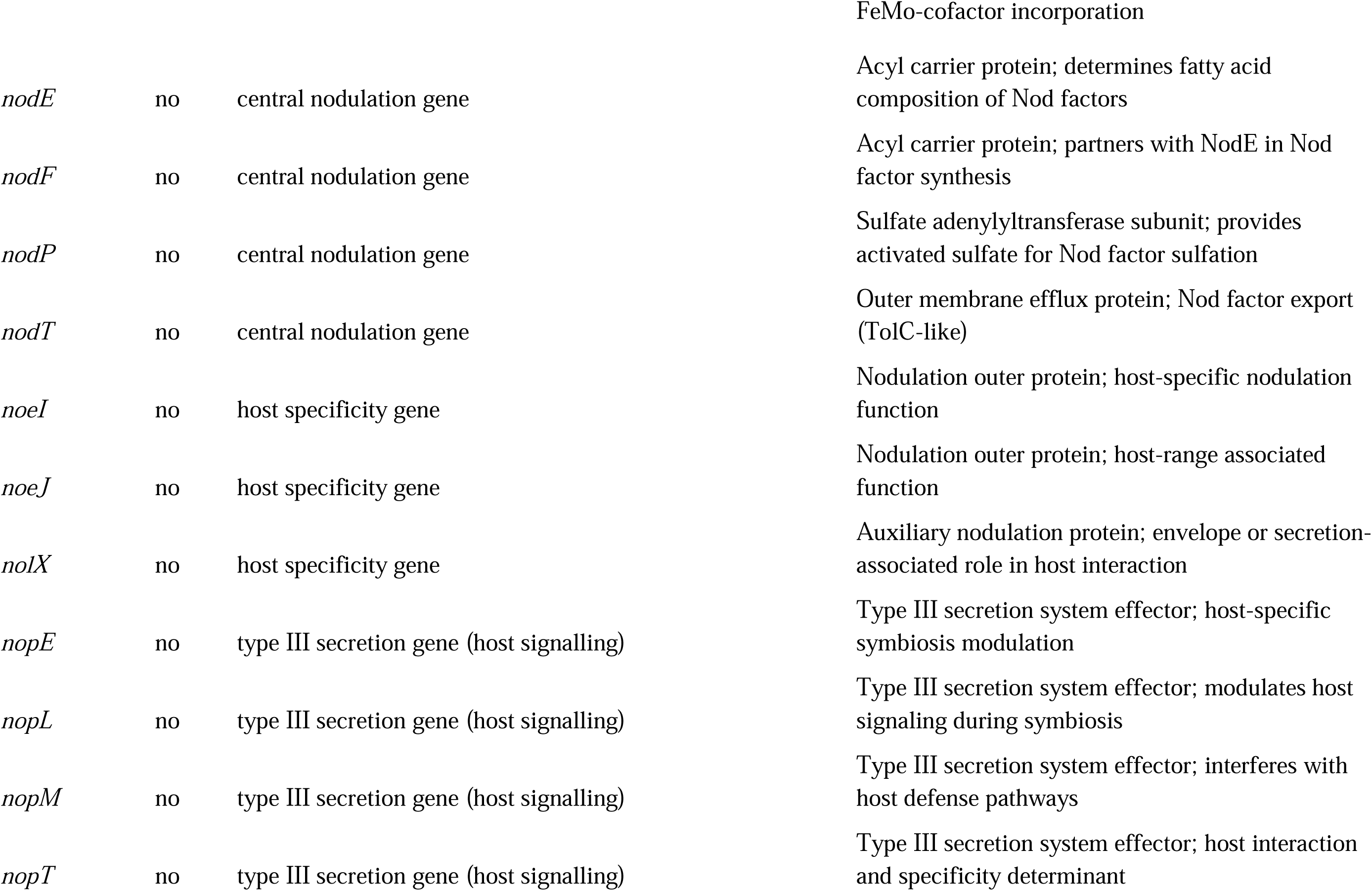
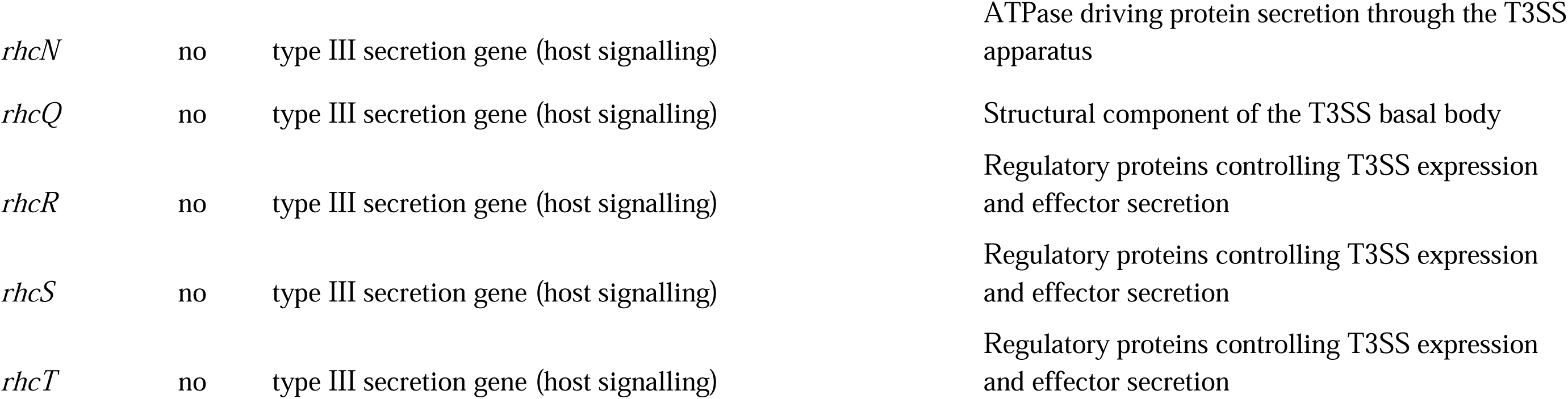
List of symbiosis island genes (see Fig. 7), with descriptions of biological role. Most but not all genes plotted in Fig. 7 were in the sequence capture kit; those that were missing are mostly organism-specific genes and are marked “no” in the second column below.

